# Complex Systems Analysis Informs on the Spread of COVID-19

**DOI:** 10.1101/2021.01.06.425544

**Authors:** Xia Wang, Dorcas Washington, Georg F. Weber

## Abstract

The non-linear progression of new infection numbers in a pandemic poses challenges to the evaluation of its management. The tools of complex systems research may aid in attaining information that would be difficult to extract with other means. To study the COVID-19 pandemic, we utilize the reported new cases per day for the globe, nine countries and six US states through October 2020. Fourier and univariate wavelet analyses inform on periodicity and extent of change. Evaluating time-lagged data sets of various lag lengths, we find that the autocorrelation function, average mutual information and box counting dimension represent good quantitative readouts for the progression of new infections. Bivariate wavelet analysis and return plots give indications of containment versus exacerbation. Homogeneity or heterogeneity in the population response, uptick versus suppression, and worsening or improving trends are discernible, in part by plotting various time lags in three dimensions. The analysis of epidemic or pandemic progression with the techniques available for observed (noisy) complex data can aid decision making in the public health response.

## Introduction

The spread of infectious diseases depends on pathogen factors (virulence), host factors (immunity), and – on the population level – on countermeasures taken by the community. Such measures cover a broad spectrum of possible engagements, and they may be highly consequential for the course of an epidemic or a pandemic [1]. The analysis of acute infectious progression in a society is critical for gauging the effectiveness of public health responses, but it is made difficult through the non-linear nature of the underlying process. Conventional approaches of reductionist research or common linearization techniques are not meaningfully applicable.

Various strategies have been employed to account for the complexity of infectious propagation. The spread of COVID-19 has been modeled with machine learning [2], networks of compartments [3] and cellular automata [4]. Power laws have been inferred [5]. Such investigations are of value, even though they are inevitably based on idealizing assumptions. In addition to modeling approaches, the analysis of actually observed data is of critical importance. The numbers in such data sets are noisy, and they are eminently non-linear (also described as “complex data” or “observed chaotic data” [6]). Complex systems research has made techniques and algorithms available to extract information from observed non-linear data series.

The manifestations of the COVID-19 pandemic have varied widely among geographic areas, when compared across countries [3,7,8] as well as across US states [9], depending on when the virus reached them, what the population characteristics were at the time of onset, and what actions were taken in response to the infectious spread. Here, we set out to investigate underlying patterns. We apply basic tools of complex systems research to compare the spread of COVID-19 in distinct countries, characterized by their varying approaches to the pandemic, from its beginning stages through early or late October 2020. Further, we compare various regions within the USA, which has left major decisions to the individual states. Patterns are discernible in Fourier and wavelet analyses. Order can be detected in time-lagged plots. Therefrom, quantitative measurements are obtainable, including autocorrelation, average mutual information, fractal dimension, and embedding dimension, which inform on the pandemic progression.

## Methods

### Source data

Here we analyze the new infections per day, either as absolute numbers or as rates per 10,000 inhabitants. The source data utilized for the present analysis came from Bing COVID-19 Tracker (www.bing.com/covid).

### Fourier spectrum and univariate wavelet analysis

Fourier analysis evaluates the spectral density by relative numbers of new infections (case rates per 10,000 inhabitants) versus frequency or versus period. Wavelet analysis does not assume stationarity in the time-series. Thus, it allows the study of localized periodic behavior. In particular, we look for regions of high-power in the frequency-time plot. The calculations for wavelet analyses of new infections were done in R. In WaveletComp, the null hypothesis, that there is no periodicity in the series, is tested via p-values obtained from simulation, where the model to be simulated can be chosen from a range of options [10]. The algorithm analyzes the frequency structure of uni- or bivariate time series using the Morlet wavelet. The time series to be analyzed was standardized, after detrending, in order to obtain a measure of the wavelet power, which is relative to unit-variance white noise and directly comparable to results of other time series. Detrending is accomplished using polynomial regression. Where indicated, all graphs are normalized to the same y-axis scale.

### Bivariate wavelet analysis

We conducted bivariate analysis of lagged data (t versus t+7 or t+14 or t+21) for joint periodicity. The concepts of cross-wavelet analysis provide tools for comparing the frequency contents by two time series as well as for drawing conclusions about their synchronicity at certain periods and across certain ranges of time. While cross-wavelet power corresponds to covariance in the time domain, wavelet coherence is a time-series measure similar to correlation. Two waves are coherent if they have a constant relative phase. The bivariate analysis results include the cross-wavelet power plot, the wavelet coherence plot, the average power plot and the phase difference image. The cross-wavelet power and coherence plot contain arrows showing the area of significant joint periods (significance level = 0.05). The direction of these arrows indicating the direction of phase differences. Up-right pointing arrows indicate that the two series are in phase and x(t) series leads, while down-right pointing arrows indicate the two series are in-phase and x(t+n) series leads. Similarly, up-left pointing arrows express that the two series are out of phase and x(t+n) series leads, while down-left pointing arrows express that the two series are out of phase and x(t) series leads. The arrows are only plotted within white contour lines indicating significance at the 10% level. A more explicit global view of the phase difference can be produced with (π/2, π) and (−π, - π /2) for out of phase and (−π /2, π /2) for in-phase. The time-averaged cross-wavelet power provides a summarized view on the shared periods, the corresponding power and the statistical significance. Cross-wavelet plots may mark areas significant due to one series swinging widely, rather than two series sharing a joint period. To avoid this false positive readout, it is more appropriate to examine wavelet coherence plots, like the coefficient of correlation. It has a value range between 0 and 1 and it shows statistical significance only in areas where the two series actually share jointly significant periods.

### Return plots

From the total numbers of new infections, we generated return plots with increasing lags, plotting daily changes x(t+1),…,x(t+7) versus x(t) and weekly changes x(t+14),…,x(t+49) versus x(t). Short time lags tend to cluster around the 45°angle, whereas increasing time delays reveal the structure of the oscillations. When graphed in 3 dimensions, these diagrams can aid in reconstructing the underlying attractor.

### Autocorrelation

A time series sometimes repeats patterns or has other properties, whereby earlier values display some relation to later values. The autocorrelation statistic (serial correlation statistic) measures the degree of that affiliation as it refers to linear dependence. The magnitude of its dimensionless number reflects the extent of similarity. The formula for autocorrelation *R_m_* is comprised of terms for autocovariance and variance

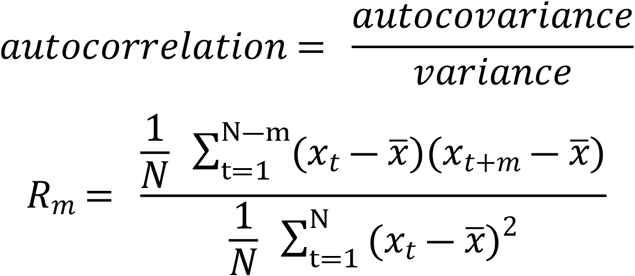

Autocorrelation coefficients range from −1 to +1, with +1 indicating perfect synchrony and −1 reflecting exact mirror images. An absence of any correlation yields *R_m_* = 0.

### Box counting dimension

The dimension of a fractal is best described as a non-integer. The dimension is a quantitative measure for the evaluation of geometric complexity by objects. A general relationship assumes

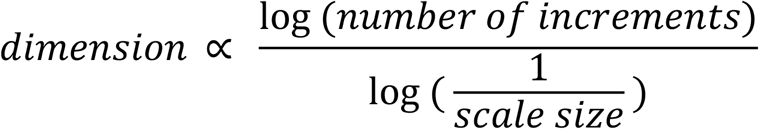

Here, the characteristic of dimension is that it specifies the rate, at which the number of increments varies with scale size. We calculated the box counting dimension after binning into 16 x 16 squares of 2-dimensional return plots with various lags.

### Average mutual information

The average mutual information (ami) represents a non-linear correlation function, which indicates how much common information is shared by the measurements of x(t) and x(t+n). The average mutual information was calculated with the mutual function R package tseriesChaos. It estimates the mutual information index for a specified number of lags. The joint probability distribution function is estimated with a simple bi-dimensional density histogram.

### Embedding dimension

Here by R package nonlinearTseries, we first use the timeLag function to decide the optimal time lag *τ* based on the average mutual information and then by the estimateEmbeddingDim function to assess the optimal embedding dimension m. Then the optimal set of regressors related to x(t) is x(t-*τ*),…,x(t-(m-1) *τ*), x(t-m*τ*).

## Results

### 1. Comparison across Countries

Across countries, a wide spectrum of measures was taken to curb the spread of SARS-CoV2. This resulted in a range of very different progression curves when graphing the numbers of new infections over time (Figure 1). India, Brazil, Sweden, Italy and the United States have been considered as hard-hit for their own internal reasons. France, Germany, over a long period Poland, and South Korea had tighter control and a less aggressive spread. All curves display close to linear ramp-up phases, followed by more or less irregular oscillations. The levels of success at suppressing the new infection rates diverged among countries, and several are experiencing a second peak.

**Figure 1:**
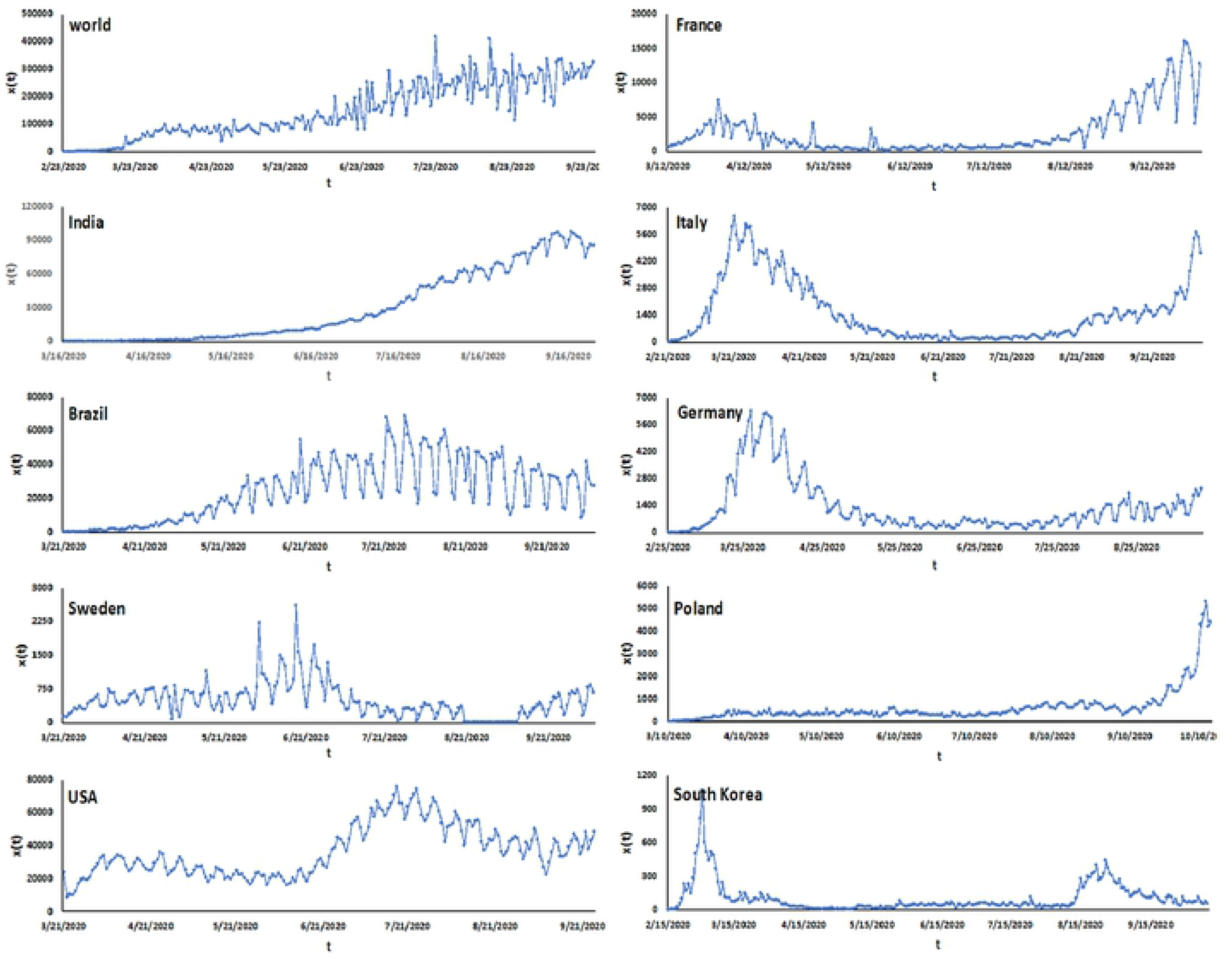
Time-Course of Disease Spread by Country. Numbers of new cases, x(t), per day versus time (t, indicating the date). Shown are the curves for (top to bottom, left to right) the globe, India, Brazil, Sweden, Italy, USA, France, Germany, Poland, and South Korea. Note the different scales of the y-axes.

Wavelet methodology aids in studying periodic phenomena in time series, particularly in the presence of potential frequency changes over time. For cross-country evaluations, all graphs were plotted on the same scale (Figure 2A). Each country was also plotted on its own scale (Figure 2B). The univariate analysis of the time course for the countries under study shows prominence of the recent upswing in France (heat intensity on the right margin of the graph). By contrast, there is comparatively more successful management by Italy, Germany, Poland and South Korea through October 2020. India, Brazil, Sweden, and the United States display cyclical fluctuations of various durations, none of which have been contained. A period of 7 days is prominent in the fluctuations of most countries, which may reflect real cyclicity or weekly reporting habits. The worldwide data are displayed in Figure S1.

**Figure 2:**
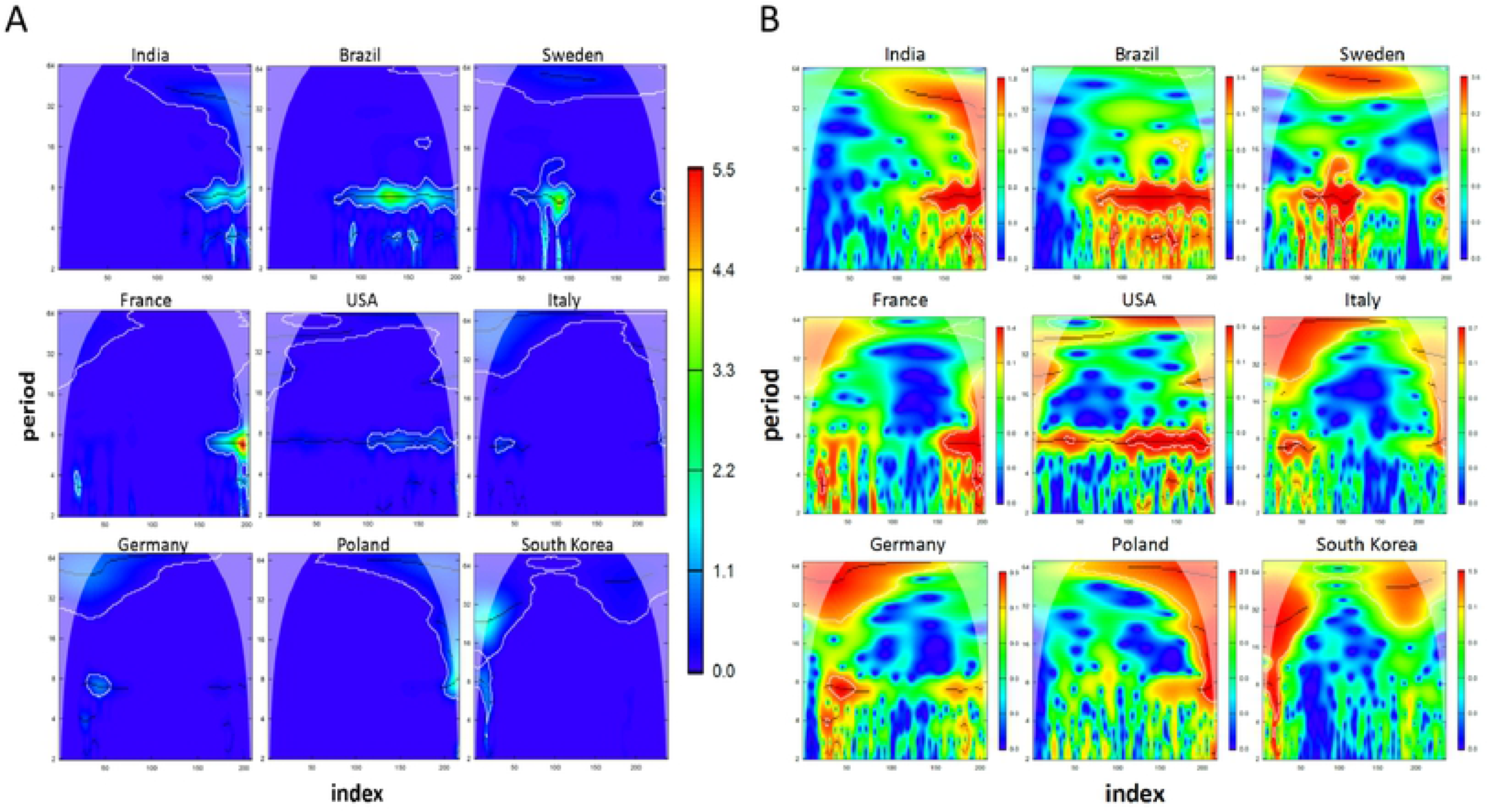
Univariate Wavelet Analysis. Cross-Wavelet Power Spectrum in the Time-Period Domain. The x-axis (index) displays the time progression, whereas the y-axis depicts the length of the period. White contour lines indicate significance of periodicity on the 0.1 level for probability of error. Lines represent the ridge of cross-wavelet power. The color bar reveals the power gradient. **A)** All countries on the same scale. **B)** Each country on its own scale.

For cross-country comparisons, we converted the new infection total numbers to new infection rates by relating them to 10,000 members of the population (Figure 3A). Similarly, complex systems can be analyzed with Fourier analysis. We first plotted Fourier power spectra versus frequency for the rates of new infections (Figure 3B). Spectral density range (high in Brazil, low in South Korea) and frequency distribution provide a readout for infectious spread. The spectral density of the normalized rates (identically scaled y-axes) (Figure 3C) confirmed good management of the pandemic spread in Germany, Poland, and South Korea (and to some degree in Italy). Despite the progressive increase in the numbers of infections in India, on a population basis, control has apparently not been lost through October 2020. By contrast, the power spectra for Brazil, Sweden, and France are reflective of potentially adverse developments. The United States display an anomaly with a periodic behavior that has a prominent cycle around 100 days.

**Figure 3:**
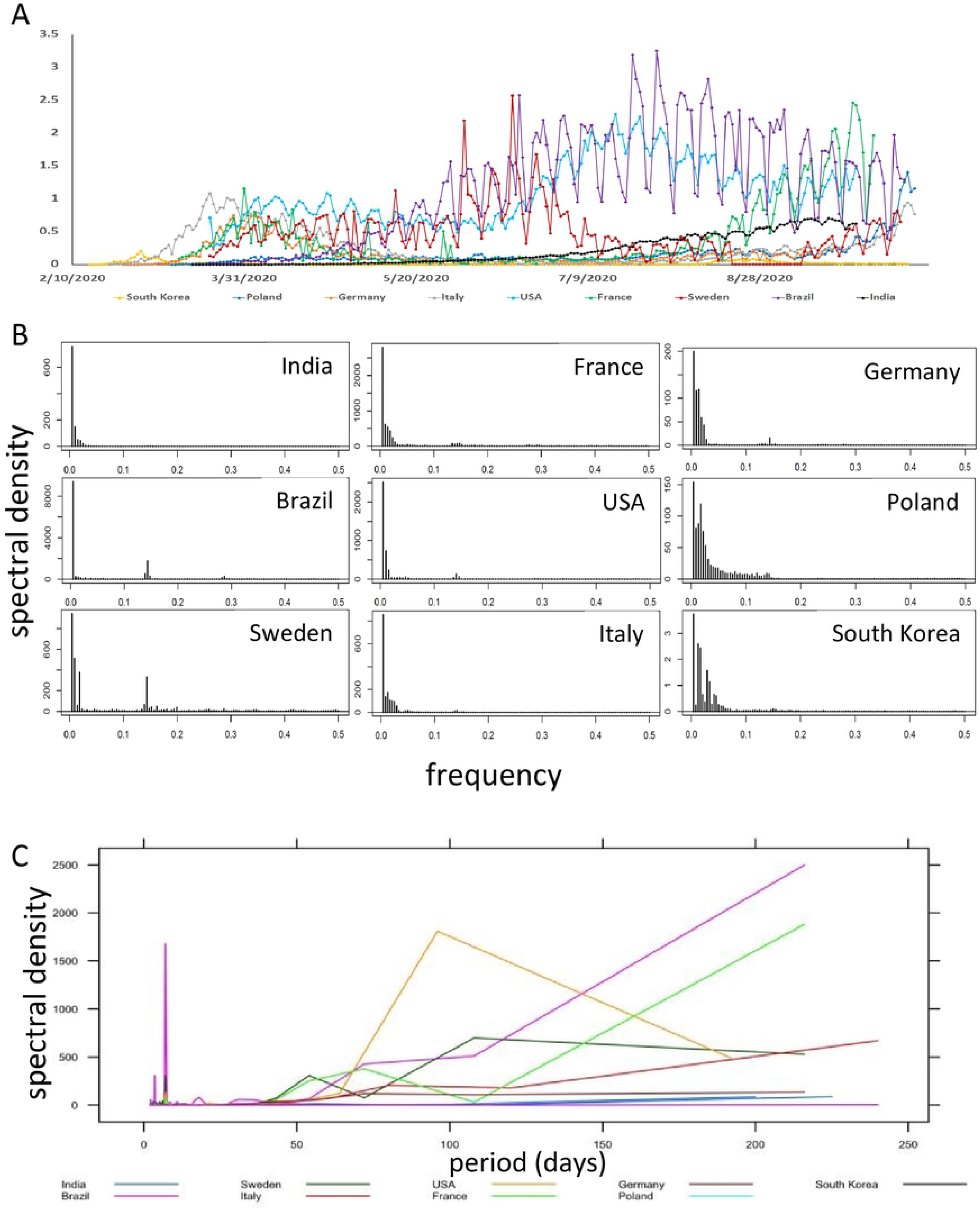
Fourier Analysis. **A) New Infection Rates.** Daily reported new numbers of infections divided by 10,000 inhabitants. The x-axis shows the calendar date. **B) Power Spectrum.** Fourier power spectra versus frequency for. new infections per 10,000 inhabitants per day in each of 9 countries. **C) Normalized Power Spectrum.** Spectral density (y-axis) versus period (in days) for infection rates per 10,000 inhabitants (x-axis). The curve shows the smoothed spectral density estimates. All y-axes have the same scale.

To gain a better understanding of the dynamics, with which disease spread occurs, we investigated progressive numbers of new infections in comparison to their increasing time lags. This approach may reveal periodicities or aid in the visualization of attractors. Expectedly, short time delays were associated with little change. With a lag time of about 7 days onward, distinct patterns emerged among countries. According to bivariate wavelet analysis for time-delayed data series (including the cross-wavelet power plot, wavelet coherence plot, average power plot and phase difference image), Italy, Germany and South Korea shared significantly joint periods of 1-2 months in the comparison x(t) versus x(t+7). South Korea has comparatively high power and significant shared periods around 3 weeks at the early stage and later the significant shared periods are also 1-2 months. The remaining countries all have segments of shorter periods (around 7 days) and longer periods shared. For India, Brazil, France, USA and Poland, the shared 7-day period only appear significant in the later part of the series. Similar results are observed in the analyses for x(t) versus x(t+14) and x(t) versus x(t+21). The phase difference plots show that in the shared longer periods, x(t) are mostly in phase with x(t+7), while they gradually become out of phase in x(t) versus x(t+14) and x(t) versus x(t+21), thus making longer lags more discriminating and informative (Figure 4A and Figure S2A,B). A reduction in cross-wavelet power levels is apparent in Italy, Germany and South Korea. Poland and France are experiencing recent increases. India, Brazil and the USA have had protracted periods of high cross-wavelet power levels. Containment is associated with longer periodicity in the distribution of cross-wavelet power. This is the case for South Korea, Germany and Italy. High cross-wavelet power around a periodicity of 7 days is reflective of poor control.

**Figure 4:**
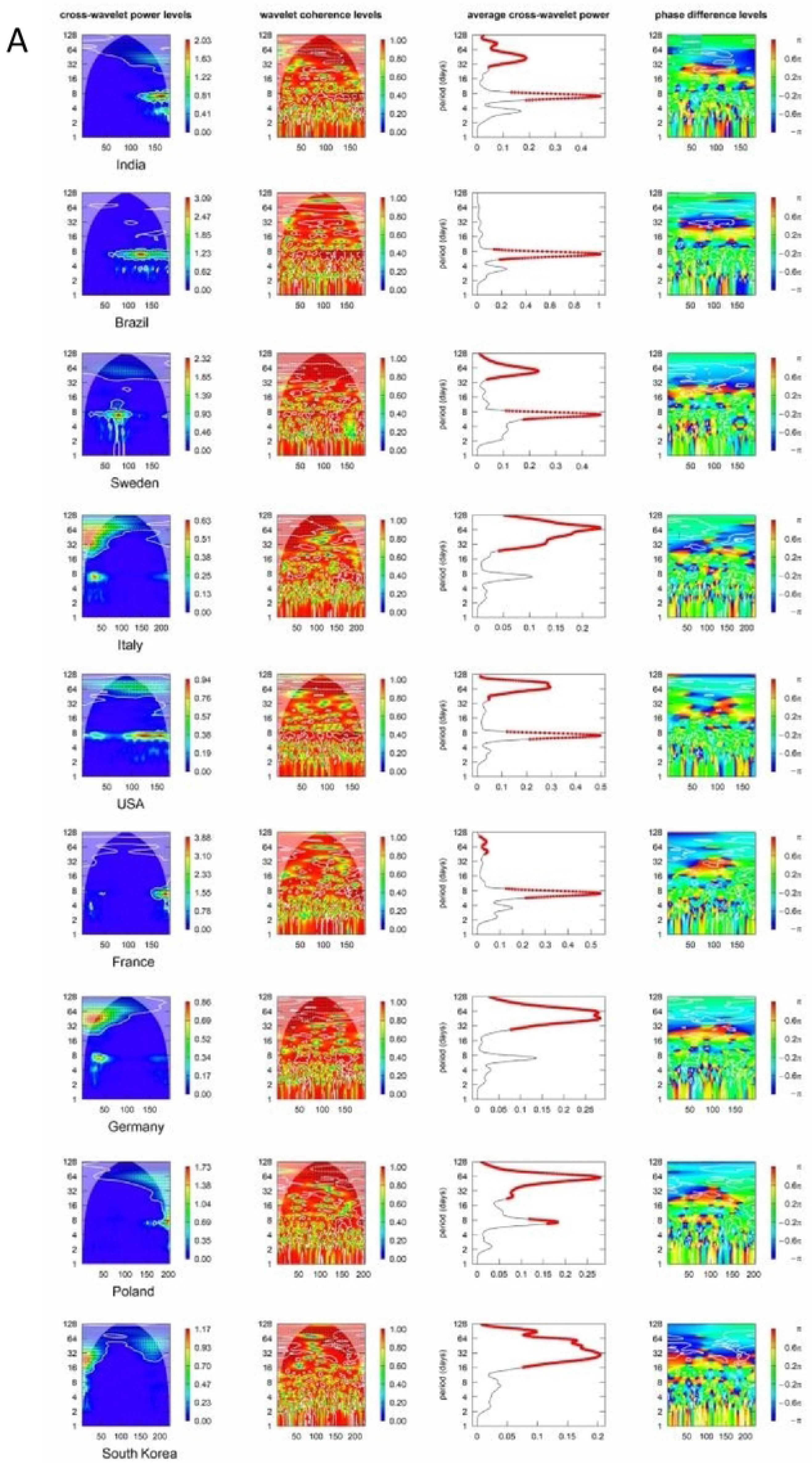
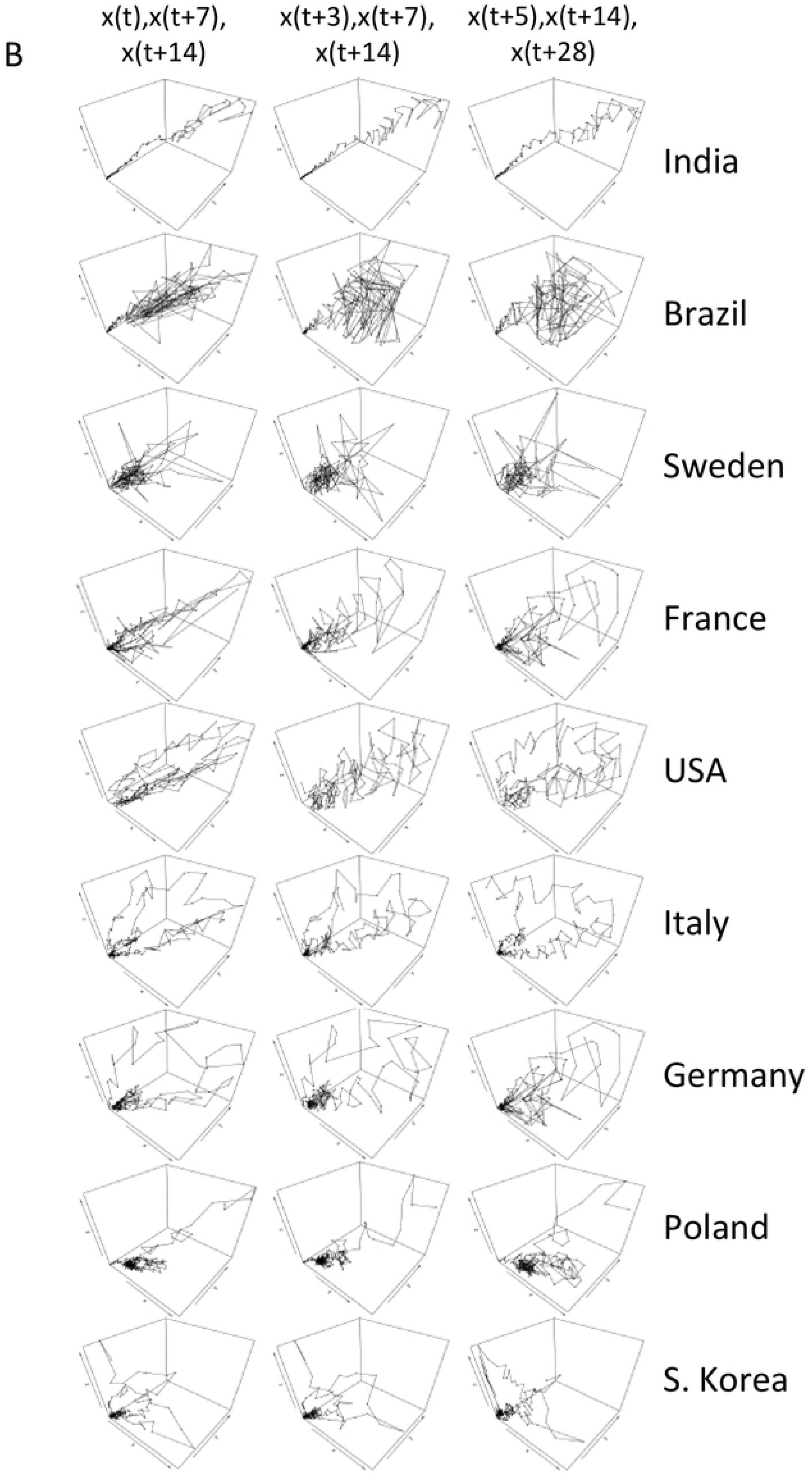
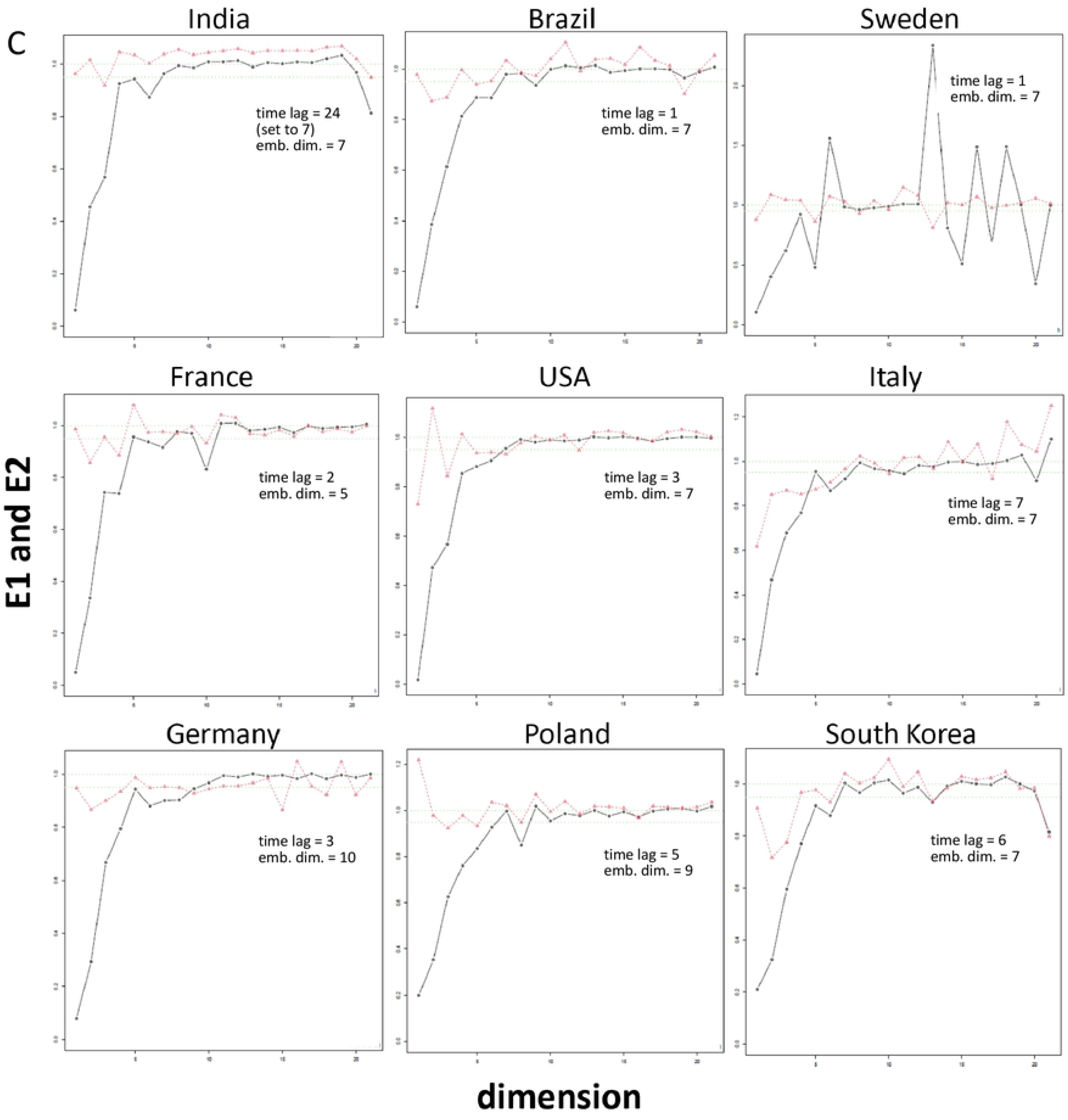
Time-Lagged Data Analysis. **A) Bivariate Wavelet Analysis**. Shown are cross-wavelet power plot, wavelet coherence plot, average power plot and phase difference image (from left to right in each row) Time-lagged data were used for x(t)/x(t+14) (for the lags x(t)/x(t+7) and x(t)/x(t21) see Figure S2). White contour lines indicate significance for joint periodicity, black arrows depict the phase difference in the areas with significant joint periods. The solid red dots on the average power plot (the third from the left) depict significant joint periods at a probability of error of 0.1. Where shown, the color bars reveal the ranges of cross-wavelet power levels. **B) Return Plots in 3 Dimensions.** Time-lagged return plots in 3 dimensions are shown, from left to right, for x(t)/x(t+7)/x(t+14), x(t+3)/x(t+7)/x(t+14), and x(t+5)/x(t+14)/x(t+28). Each country of interest has its own row. **C) Embedding Dimension.** The plots show how Cao’s algorithm uses 2 functions in order to estimate the embedding dimension from the time series (the E1(d) and E2(d) functions), where d denotes the dimension.

To generate informative return plots, we utilized 3 dimensions, which allows for the visualization of two lags from x(t) (or a from a later start point) and may reveal the pattern of an attractor. In this depiction, a rapid increase or decrease in new infections is reflected in a close-to straight line, oscillations generate a near-toroid attractor, while successful management shrinks the torus and moves it closer to the origin. Initially, we evaluated multiple time delays. Most discriminating were x(t)/x(t+7)/x(t+14), x(t+3)/x(t+7)/x(t+14), and x(t+5)/x(t+14)/x(t+28) (Figure 4B). The progressive increase in new cases over the time period in India is reflected in a predominantly linear curve on each scale. The wide fluctuations in Brazil generate a largely disordered appearance. Disorder is also apparent in Sweden. France initially managed the pandemic well, but is experiencing a dramatic upswing, which obscures order. Cyclical patterns, suggesting the outlines of attractors, are apparent in USA, Italy, Germany, and South Korea (where most data points are concentrated near the origin). Poland initially displayed a well-contained attractor, but the recent substantial upswing in new infections is reflected in a linear progression from there (for separate analyses of the two phases, see Figure S3). We also calculated the embedding dimensions for the lagged data (Figure 4C). Germany has the highest embedding dimension of 10, followed by Poland with 9. Several countries have an embedding dimension of 7, including Brazil, Sweden, USA and South Korea. Italy and France have the embedding dimension equal to 5. India is unusual due to its longer lag period of 24 days. When the lag period is set at 7 days, the embedding dimension of India is also equal to 7. For the worldwide data, the calculated embedding dimension is 7 with a time lag of 1 (not shown).

The autocorrelation of two data strings with short time lags is expected to be high (approaching 1.0) because there is little opportunity for dramatic change (high infection rates on day t likely produce similarly high numbers on the consecutive day t+1, while low numbers are followed by few new infections on the next day). Autocorrelation may remain high for extended lags in the initial ramp-up and at the oscillatory stage, depending on the regularity of the fluctuations. A society that succeeds in curbing the disease spread will leave the highly correlated initial ramp-up and consecutive oscillatory phases, thus displaying a gradual decrease in values at the longer lags. The decline in the autocorrelation numbers of progressively lagged data by country appeared to be reflective of the stringency, with which the pandemic was addressed (Figure 5A). From a lag of 6 onward, Poland and South Korea have substantially declining values (although due to the recent steep upswing in new infections, Poland deviates from the trend at very long lags), Germany shows a dramatic lowering at a lag of 21 and above. By contrast, India and Brazil stay uniformly high. So do the global numbers, which are inherently heterogeneous.

**Figure 5:**
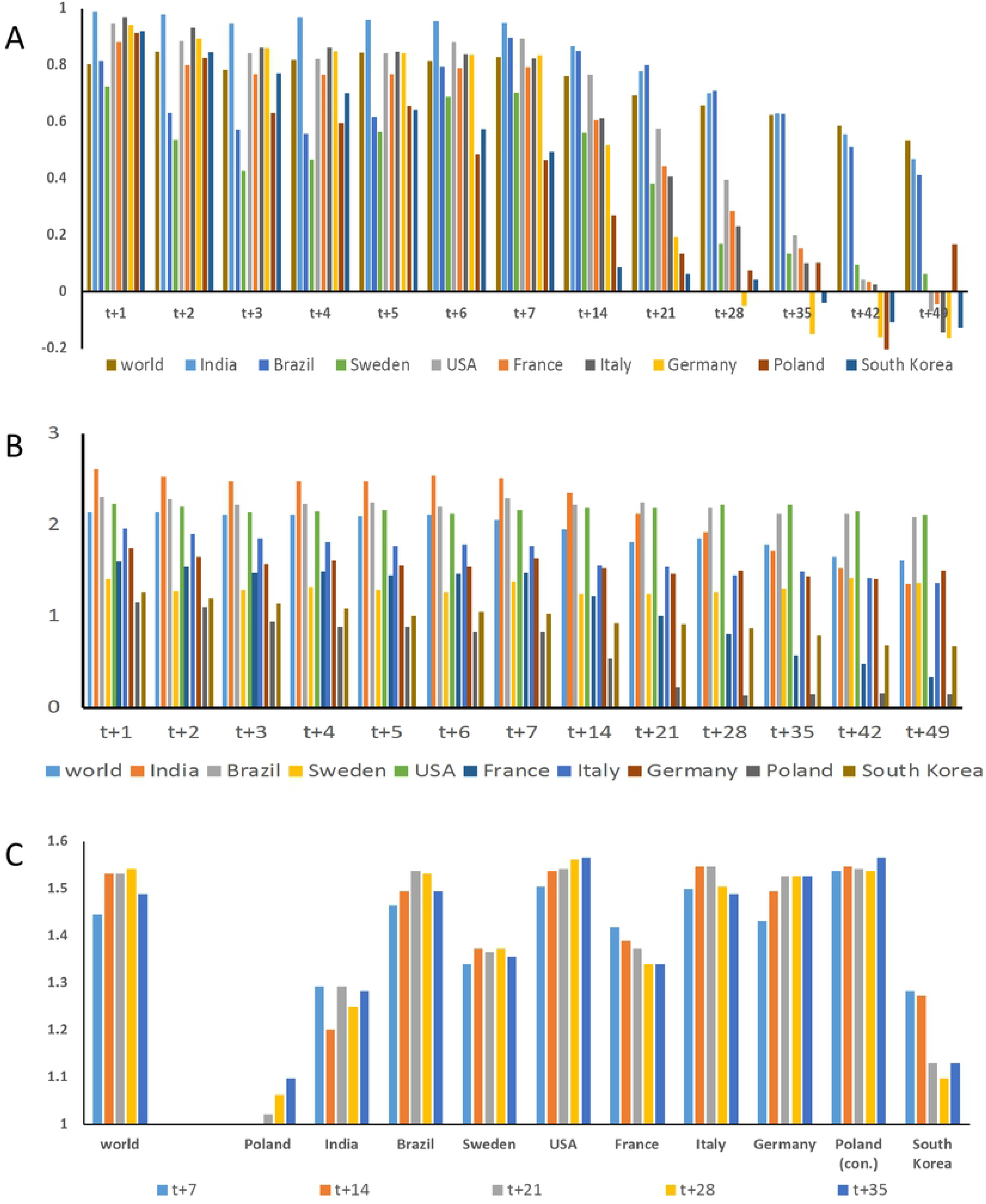
Readouts of Complexity for Lagged Data on COVID-19 Spread by Country. **A) Autocorrelation.** Bar graph of the autocorrelation in COVID-19 spread with each bar color representing a different country. The selected time lags are indicated on the x-axis, all are calculated versus x(t). **B) Average Mutual Information.** Bar graph of average mutual information in COVID-19 spread with each bar color representing a different country. The selected time lags as indicated on the x-axis are all calculated versus x(t). **C) Fractal Dimensions.** Box counting dimensions are calculated for 2-dimensional return plots of increasing lags, x(t+7) versus x(t) through x(t+35) versus x(t). 9 countries are evaluated, and the worldwide numbers are shown on the left. Poland is represented twice, over the entire evaluation period through 12 October 2020 (which contains a steep incline) and over the shorter phase of containment through 18 September 2020 (cont. = contained period).

The average mutual information reflects information shared by the measurements of x(t) and x(t+n). Expectedly, it declines with increasing lag. Poland starts with a relatively low value (1.15 at t versus t+1) and shows a rapid decrease with longer lag. It then stays around at a low level of 0.15 from lags of 21 to 49 days. France displays a gradually decreasing trend with the average mutual information starting at 1.60 and ending at 0.34 at the lag of 49 days. India shows a similar pattern as France but with much higher average mutual information (due to the constant uptick in numbers), ranging between 2.61 and 1.37. Four other countries, including Germany, USA, Sweden and Brazil, all express relatively flat average mutual information values, staying around levels of 2.20 for the USA and Brazil, 1.5 for Germany, and 1.3 for Sweden. Reflecting progressively improved control, Italy and South Korea also have decreasing trends, but much flatter at 1.96-1.36 for Italy and 1.26 to 0.66 for South Korea, respectively (Figure 5B).

A rapid increase in new infections is reflected in a small fractal dimension (practically approximated by the box counting dimension with values between 1 and 2) of the 2-dimensional return plots with progressive lags. Intermediate phases are characterized by higher fractal dimensions (approaching 2), depending on the nature of the oscillations. Conversely, successful management through the reduction in new infections should be reflected in a contraction of the attractor on the return plot, which is assessable through the box counting dimension. A trend is displayed in the comparisons from shorter to longer lag periods. Distinct management strategies across different countries generate a heterogeneous pattern worldwide, rendering the fractal dimension high regardless of the lag in x(t+n) versus x(t) plots. Steep increases in new infections (Poland, India) have dimensions close to 1. Intermediate phases are characterized by higher numbers. Successful fights against the pandemic (South Korea) are causative for declining size dimensions with increasing lag (Figure 5C).

### 2. Comparison across US States

Within the USA, individual states have encountered a rather wide range of progression phenotypes in the spread of new COVID-19 infections (Figure 6). This is due to variations in international connectedness and population density (reflected in the early peaks in the Northeastern states New York and Massachusetts), holiday travel (Florida), policy decisions and other factors.

**Figure 6:**
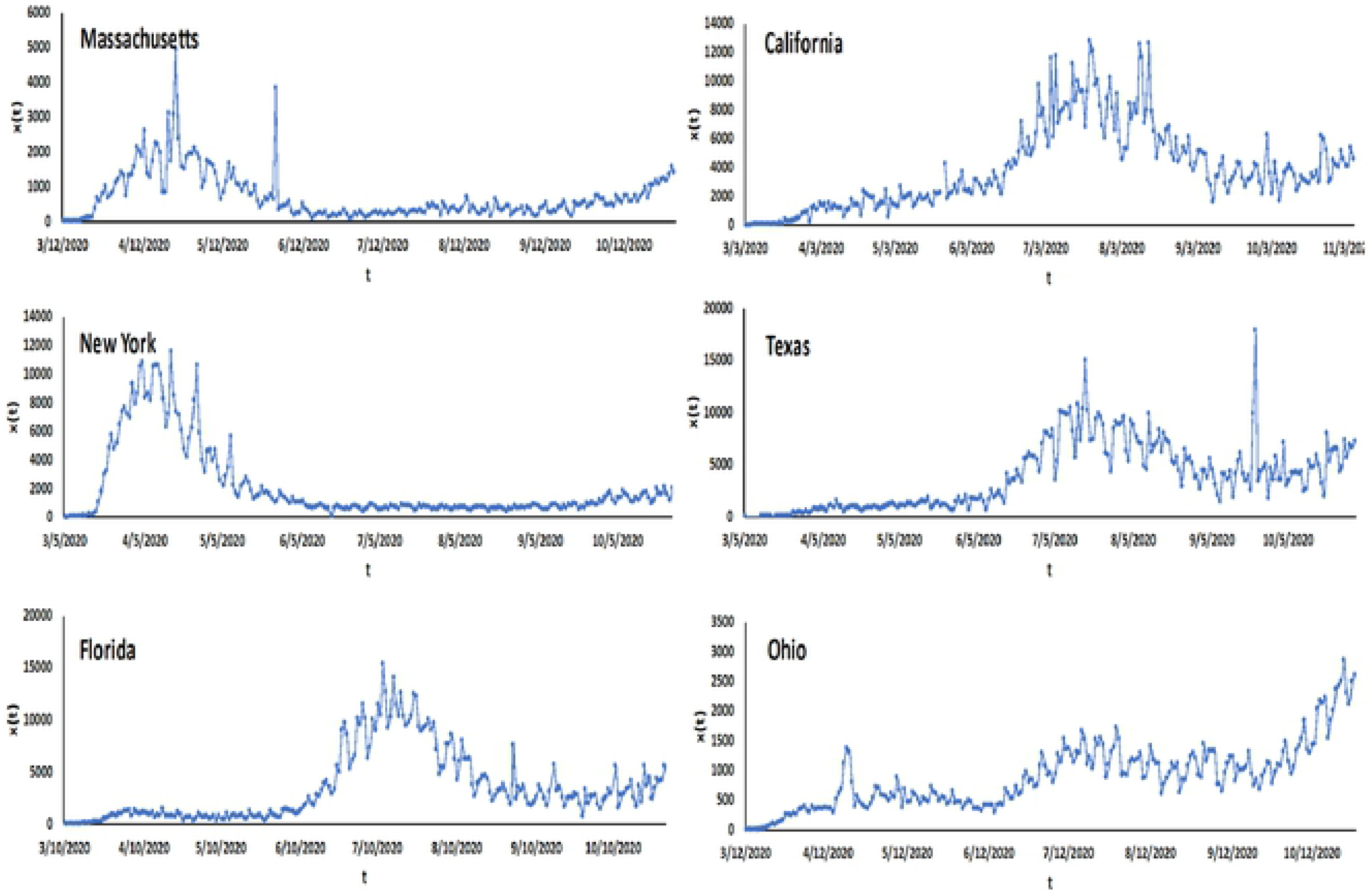
Time-Course of Disease Spread for Individual US States. Numbers of new cases, x(t), per day versus time (t, indicating the date). Shown are the curves for (top to bottom, left to right) Massachusetts, New York, Florida, Texas, California, and Ohio.

Wavelet analysis of new infections (one scale across all states) shows good control (right side of the graph) after initial affliction (left area) for Massachusetts and New York, which having had early spikes in new infections have achieved good success in containment. Through the observation period, control has not been maintained in Ohio. The periodicity in individual states (each on their own scales) is poorly defined, except for Florida and Ohio, where 7 days yield a prominent signal (Figure 7A,B).

**Figure 7:**
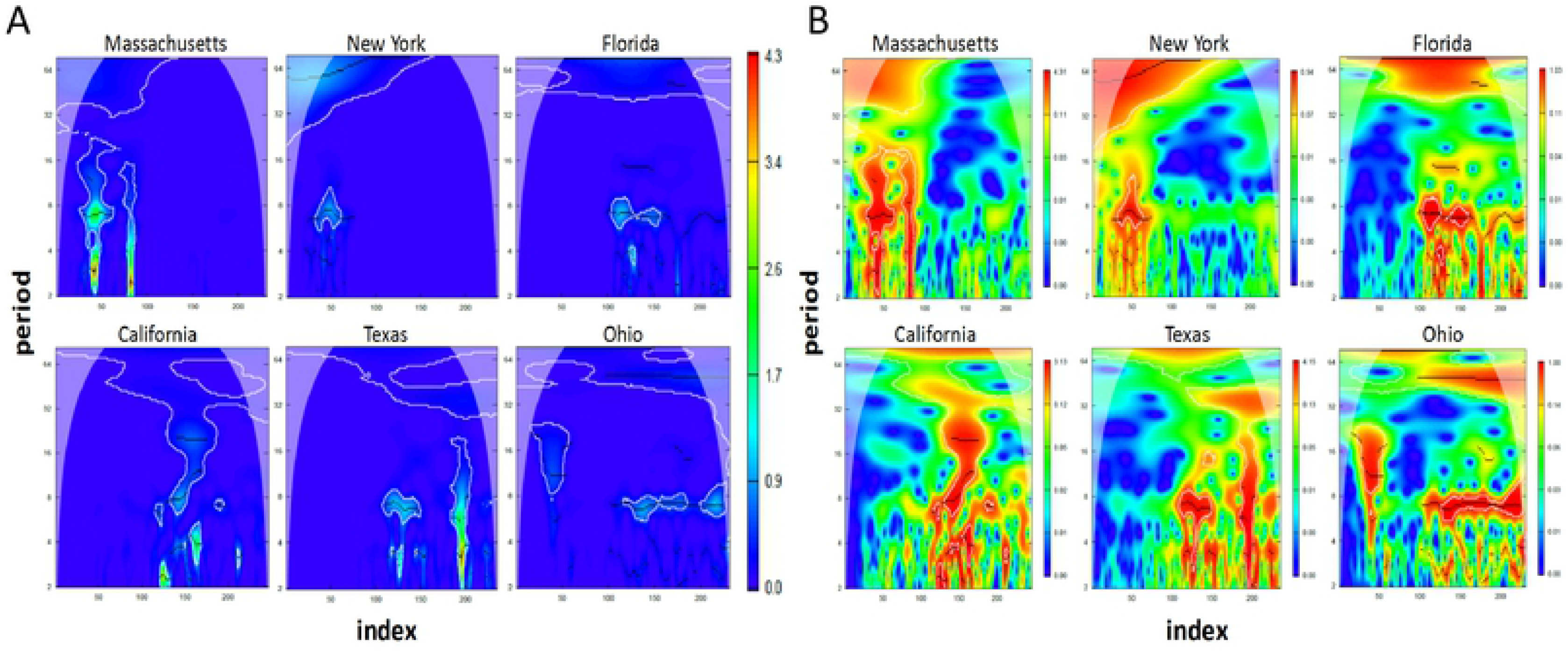
Univariate Wavelet Analysis. Wavelet power spectrum in the time-period domain. Contour lines indicate significance of periodicity with 0.1 significance level. Black lines indicate the ridge of wavelet power. The color bar reveals the power gradient. **A)** All states on the same scale. **B)** Each state on its own scale.

We normalized the new infection numbers to rates by relating them per 10,000 inhabitants (Figure 8A). Figure 8B shows the periodogram for the 6 states under investigation with frequencies between 0 and 0.10 (the graph is almost flat for the higher frequencies). There exist clear heterogeneous patterns in the comparison among these states. New York and Massachusetts display steadily decreasing spectral density values from the longest period to around 1-2 weeks (corresponding to a frequency range around 0.07-0.14). Florida and Texas share similar patterns with a few low spikes in their periodograms after the first 3 highest ones. The graph for California flattens out after the lowest three frequencies, with the longest period (the whole series) having the highest value. Ohio’s pattern is quite unique with fluctuating values from the longest periods through around 5-6 weeks. The Fourier power spectrum for the infection rates (Figure 8C) indicates similar periodic patterns as in the periodograms of Figure 8B. These patterns are less prominent due to the adjustment to the same y-axis scale (the scale reflects the magnitude of the positive rates, the shape shows the evolution of the disease).

**Figure 8:**
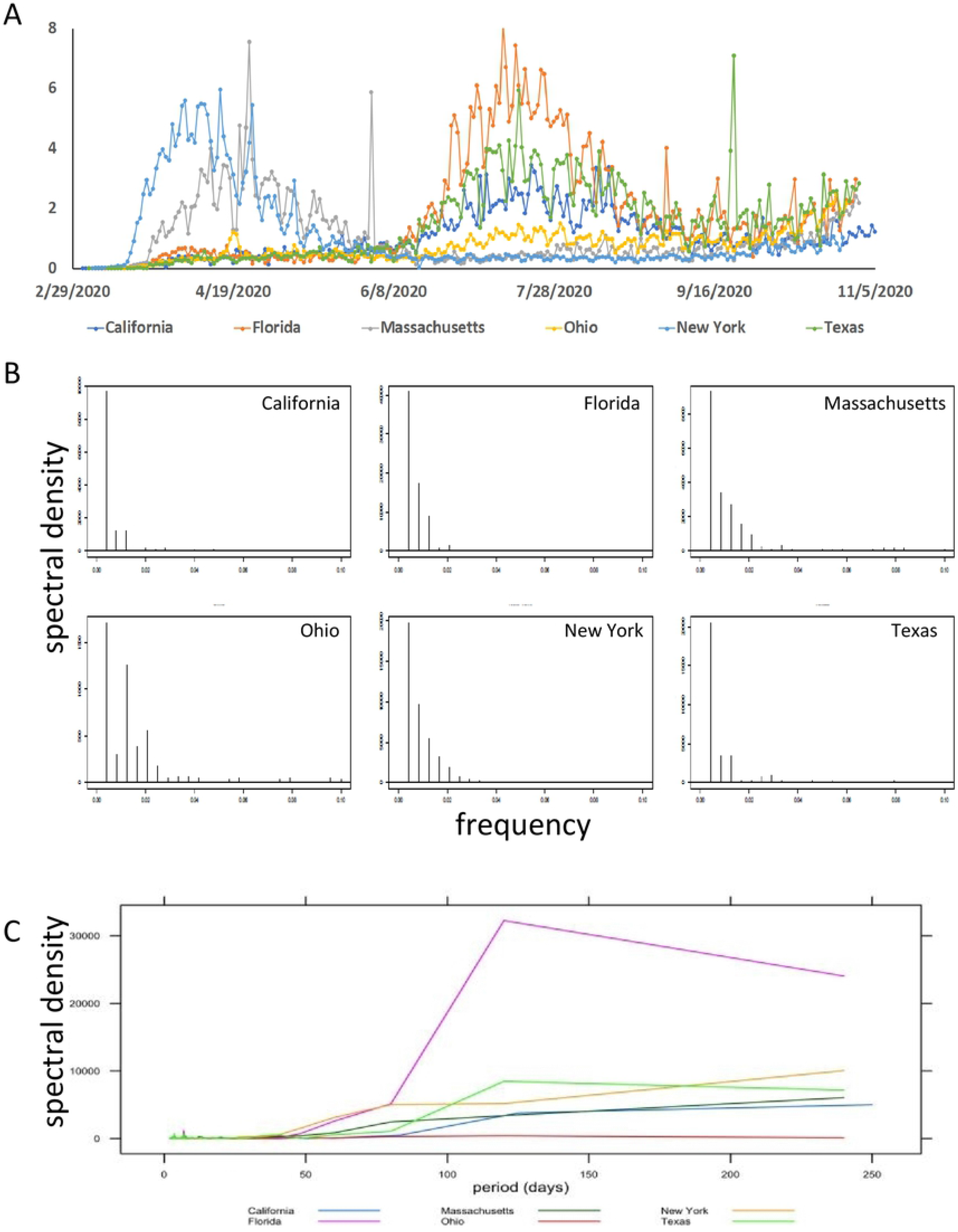
Fourier Analysis. **A) New infection Rates.** Daily reported new numbers of infections divided by 10,000 inhabitants (infection rates). The x-axis shows the calendar date. **B) Power Spectrum.** Periodogram plot on the series of the new infection rates. The x-axis is the frequency (per day) and the y-axis represents the spectral density. The y-axis ranges vary among graphs. **C) Normalized Power Spectrum.** Spectral density versus period (in days) for infection rates. All y-axes have the same scale.

We conducted bivariate wavelet analysis on the time-lagged data (Figure 9A and Figure S4). The shared synchronicity segments between x(t) and x(t+n) can be grouped into shorter periods (around 7 days) and longer periods (approximately 3 weeks, 1 month, 2 months). New York does not display substantial joint short periods. Ohio and Texas mainly have correlation at the end of the series around the 7-day period. Massachusetts experiences joint periodicity around the 7-day period at the early stage of the series. Florida and California have joint periods in the middle of the observation time frame. The levels of average cross-wavelet power are higher in states with poor control (x-axes scales for Florida, Ohio). The peak power shifts toward higher periodicity with improved control (y-axes scales for New York, Massachusetts). The return plots in 3 dimensions, utilizing the same time lags as for the countries, seemed to reflect contraction of the attractor in Massachusetts, cyclicity in New York, Florida and California, no containment in Texas, and an ejecting diagonal in Ohio which may reflect exacerbation (Figure 9B). The embedding dimensions varies among states, such that the most contained states (New York, Massachusetts) have the lowest embedding dimension (Table 1).

**Table 1:** Embedding Dimension for Time-Lagged Data by U.S. State. Embedding dimensions were calculated according to Cao’s algorithm, which uses 2 functions in order to estimate the embedding dimension from the time series. The table shows the calculated time lags and embedding dimensions for each U.S. state under study.

| state | California | Ohio | Florida | Texas | New York | Massachusetts |
| --- | --- | --- | --- | --- | --- | --- |
| time lag | 11 | 3 | 3 | 3 | 10 | 7 |
| embedding dimension | 8 | 8 | 7 | 7 | 6 | 6 |

**Figure 9:**
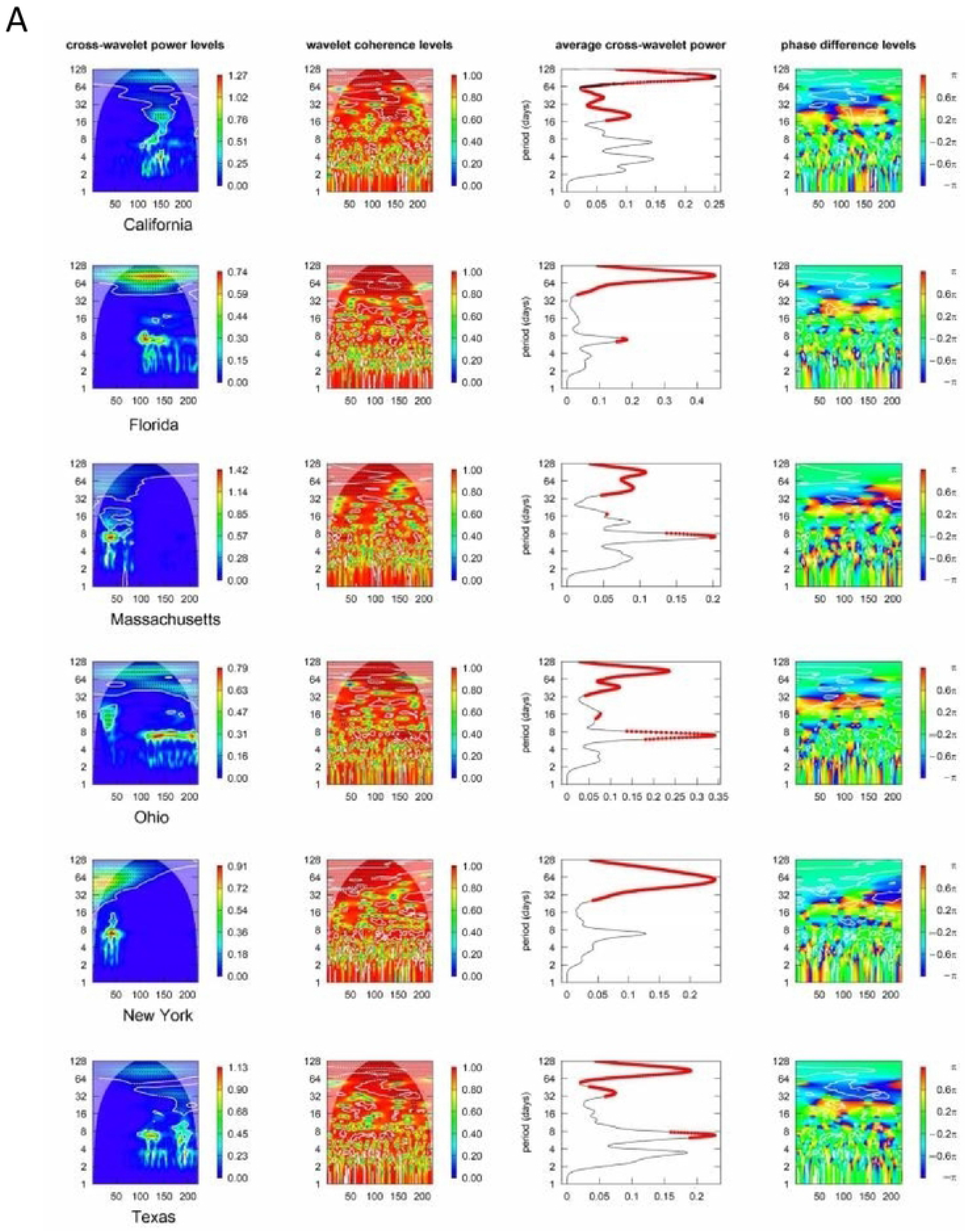
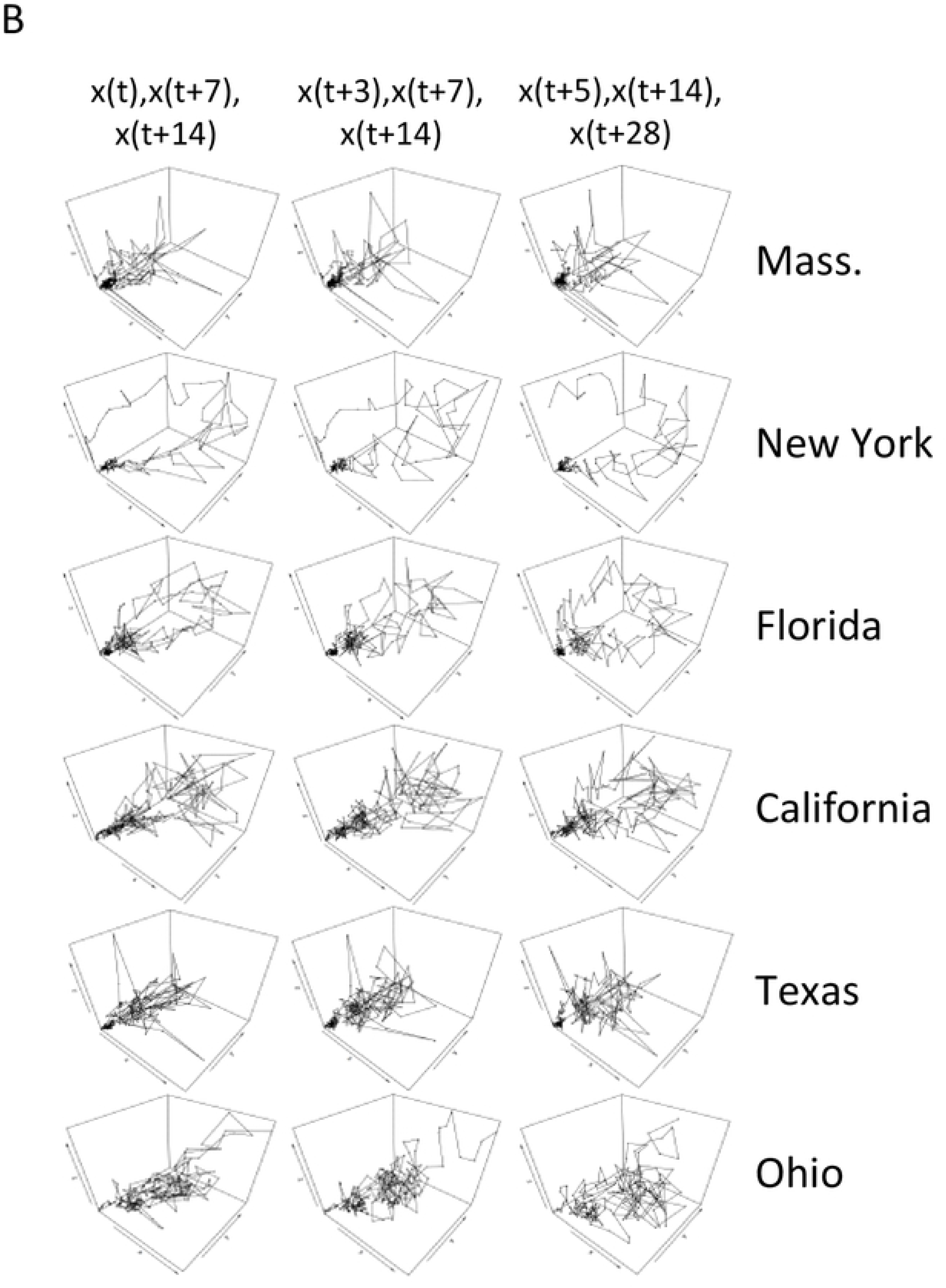
Time-Lagged Data Analysis by US State. **A) Bivariate Wavelet Analysis**. Shown are cross-wavelet power plot, wavelet coherence plot, average power plot and phase difference image (from left to right on each row) Time-lagged data were used for x(t)/x(t+14) (for the lags x(t)/x(t+7) and x(t)/x(t21) see Figure S4). White the contour lines indicate significance of joint periodicity, black arrows indicate the phase difference in the areas with significant joint periods. The solid red dots on the average power plot (the third from the left) reflect significant joint periods at a significance level of 0.1. **B) Return Plots in 3 Dimensions.** Time-lagged return plots in 3 dimensions are shown, from left to right, for x(t)/x(t+7)/x(t+14), x(t+3)/x(t+7)/x(t+14), and x(t+5)/x(t+14)/x(t+28). Each state under investigation has its own row.

The autocorrelation for return plots of increasing lags show a progressive decline in the numbers of New York and Massachusetts, which implemented strong containment measures after having been afflicted early. The values decline less steeply for Texas and California. Ohio displays an anomaly with increasing values for very long lags. The state, while not heavily afflicted on a per capita basis, never achieved containment, only a stationary level, and has since experienced another wave (Figure 10A). Up to a maximum lag of 49 days, the average mutual information for the 6 US states under study ranges between 1.0 and 2.0. Overall, all states show a slightly decreasing pattern except for California, which is relatively leveled at a value of 2.0 (Figure 10B). Unexpectedly, the box counting dimension (Figure 10C) is less discerning than it was for the evaluation across countries. This may be due to the much lower power conveyed by smaller population sizes.

**Figure 10:**
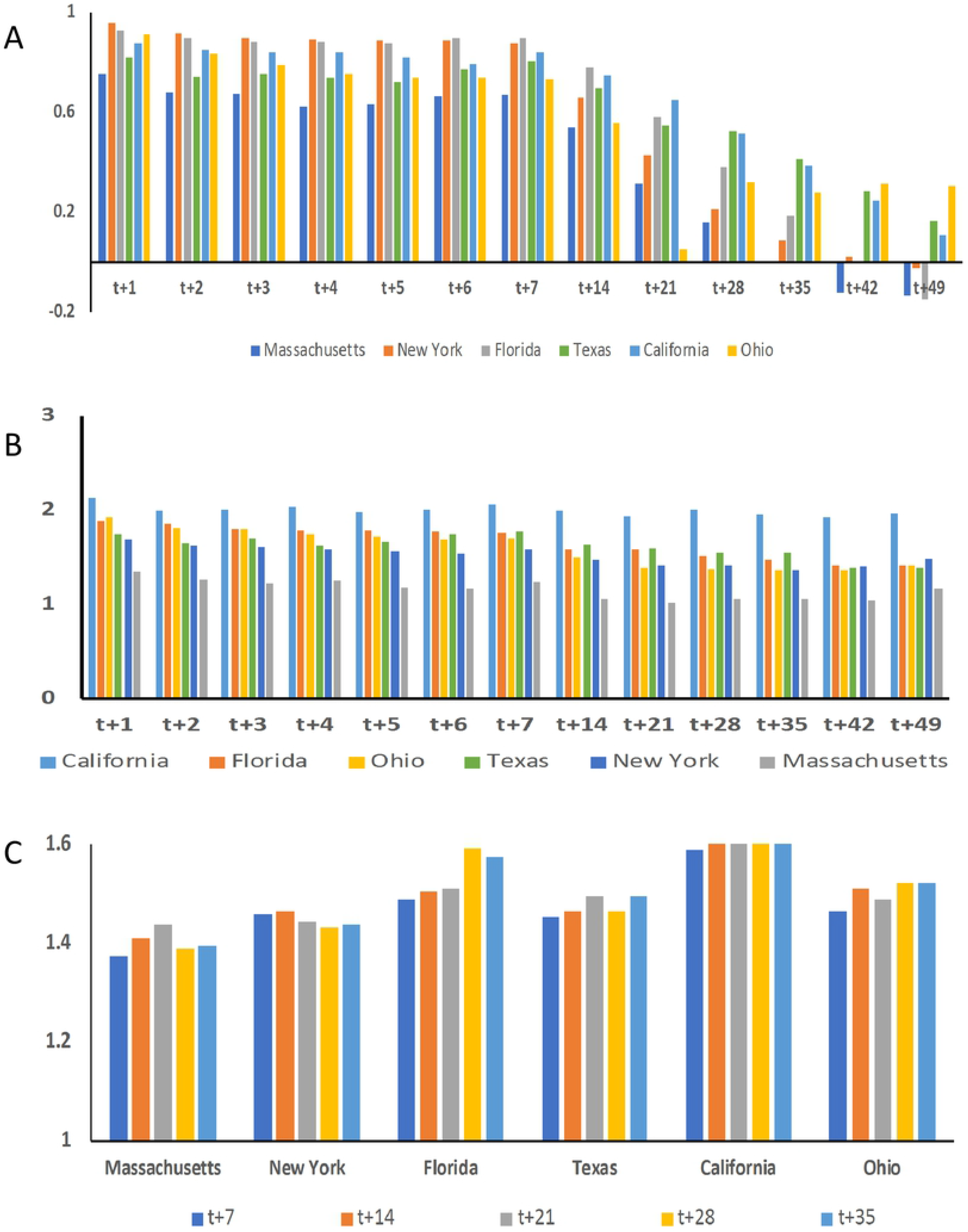
Readouts of Complexity for Time-Lagged Data by U.S. State. 6 US states have been evaluated. **A) Autocorrelation.** Bar graph of the autocorrelation in COVID-19 spread with each bar color representing a different US state. The selected time lags are indicated on the x-axis, all are calculated versus x(t). **B) Average Mutual Information.** Bar graph of average mutual information in COVID-19 spread with each bar color representing a different state. The selected time lags are indicated on the x-axis, all are calculated versus x(t). **C) Fractal Dimensions.** Box counting dimensions are calculated for 2-dimensional return plots of increasing lags, x(t+7) versus x(t) through x(t+35) versus x(t).

## Discussion

In the present investigation we find that the analysis tools for observed complex data can aid in the interpretation of pandemic spread across communities. Difficulties in analyzing the non-linear patters of infectious disease spread may be tamed by applying the tools of complex systems research. The approach can reveal patterns, where a simple time course of new cases does not. Further, non-linear analysis allows the study into various facets of the process, depending on whether the starting data are new cases, hospitalizations, deaths or other readouts. Maps can be generated and evaluated for their fractal dimensions [11]. The operational approximation of Lyapunov exponents may be meaningful, although they were largely uninformative for the present study (Supplemental Figure S5).

Among the countries analyzed, South Korea has had the most successful control of the pandemic spread according to low intensity in univariate wavelet analysis, low spectral density range in Fourier analysis, low spectral density of the normalized rates, a reduction in cross-wavelet power levels according to bivariate wavelet analysis and longer periodicity in the distribution of cross-wavelet power. Further, declining box counting dimensions, autocorrelation values with increasing time lag, and decreasing trends (at a low slope) in average mutual information confirm containment. Cyclical patterns in return plots, suggesting the outlines of attractors, are apparent and most data points are concentrated near the origin of the graph. Germany exhibited good management through October 2020 according to univariate wavelet analysis, spectral density in the power spectrum of the normalized rates, a reduction of cross-wavelet power levels in bivariate wavelet analysis, longer periodicity in the distribution of cross-wavelet power, a dramatic lowering of autocorrelation values at a lag of 21 and above, and relatively flat average mutual information values, staying around levels of 1.5. Cyclical patterns in return plots suggest the outlines of an attractor. Good control by Italy consecutive to the early impact and through October 2020 is reflected in low intensity and fluctuation when applying univariate wavelet analysis, in a reduction of cross-wavelet power levels for bivariate wavelet analysis of time-delayed data, longer periodicity in the distribution of cross-wavelet power, and decreasing trends (at a low slope) in average mutual information. Cyclical patterns in return plots, suggesting the outlines of an attractor, are apparent. Poland had two distinct phases. By univariate wavelet analysis and density in the power spectrum of normalized rates, there was indication of good management through October 2020. According to bivariate wavelet analysis for time-delayed data series and return plots, the recent substantial upswing in new infections is reflected, which also results in box counting dimensions close to 1. From a lag of 6 onward, Poland has substantially declining autocorrelation values, although due to the recent steep upswing in new infections, the trend reverses at very long lags. The average mutual information starts with a relatively low value (1.15 at t versus t+1) and shows a rapid decrease with longer lag, staying level from lags of 21 to 49 days. In the United States, univariate wavelet analysis displays cyclical fluctuations of various durations, none of which have been contained. According to bivariate wavelet analysis for time-delayed data series, there have been protracted periods of high cross-wavelet power levels. Cyclical patterns in return plots, suggesting the outlines of attractors, are apparent. The USA expresses relatively flat average mutual information values, staying around levels of 2.20. In France, univariate wavelet analysis of the time course shows prominence of the recent upswing (heat intensity on the right margin of the graph), the power spectrum is reflective of potentially adverse developments. The second wave of infections is apparent in bivariate wavelet analysis and in the obscured order in return plots. France displays a gradually decreasing trend of average mutual information. India expresses cyclical fluctuations of various durations in univariate wavelet analysis, none of which have been contained. On a population bases, the spectral density suggests that control has not been lost through October 2020. Bivariate wavelet analysis shows protracted periods of high cross-wavelet power levels, return plots reflect the progressive increase in new cases over the time period in a predominantly linear curve on each scale, box counting dimensions are close to 1, and autocorrelation values stay uniformly high with increasing time lag. India displays a gradually decreasing trend of average mutual information. Brazil experiences cyclical fluctuations of various durations in univariate wavelet analysis, none of which have been contained. By Fourier analysis, the spectral density range is high. The power spectrum is indicative of potentially adverse developments. According to bivariate wavelet analysis, there have been protracted periods of high cross-wavelet power levels. In return plots, the wide fluctuations generate a largely disordered appearance. The autocorrelation values stay uniformly high. Brazil expresses relatively flat average mutual information values, staying around levels of 2.20. Sweden shows cyclical fluctuations of various durations in univariate wavelet analysis, none of which have been contained. The power spectrum is reflective of potentially adverse developments. In return plots, disorder is apparent. Sweden expresses relatively flat average mutual information values.

Prima facie, the curves of new infections versus time for three Western European countries, France, Italy, and Germany, appear similar. Complex systems analysis reveals the upswing in France to be much more perilous than the increases in the curves of new infections by the other two countries. The management of infectious spread also requires improvements in the United States, Sweden and Brazil. The selection of the observation period can dramatically influence the results. Poland was initially very successful in containing the pandemic, but then experienced a substantial upswing. Analyzing these two phases individually or in conjunction yields very different data sets, which inform about distinct aspects of the infectious progression.

The fluctuations of new infections in an epidemic or a pandemic pose challenges to the evaluation whether a decline reflects true containment (“rounding the corner”) or just the calm before another wave. The readouts of non-linear systems analysis can aid in making such a distinction. A complex occurrence that experiences containment will strive toward a point attractor in phase space and move toward the origin. Such a progression is represented in a declining fractal dimension, and the transition from fluctuations (often associated with a torus attractor) toward limitation of new cases is expected to reduce the autocorrelation.

One constraint of complex systems analysis is the need for large data sets. In this regard, the availability of about 230 data points (daily new cases March through October 2020) for each geographic area in this study is somewhat low. The robustness of pertinent studies increases with larger data sets over time. Reporting errors could have a non-trivial impact, and may be reflected in the frequent occurrence of a peak at 7 days in the spectral analysis (possibly indicating weekly totals). This problem can be addressed by utilizing moving averages. The homogeneity or heterogeneity in management by the community under study determines the noise level. The worldwide numbers of new infections have a lot of background due to varying patterns across countries.

## Acknowledgements

GFW is supported by NIH grant CA224104.

## Supplement

**Figure S1: Power Spectrum and Univariate Wavelet Analysis for Worldwide New Cases. A)** Wavelet analysis and model fit (minimum power level: 0, significance level: 0.05, only coi: false, only ridge: false). **B)** Fourier analysis.

**Figure S2: Bivariate Wavelet Analysis by Country.** The graphs represent cross-wavelet power plot, wavelet coherence plot, average power plot and phase difference image (from left to right on each row) Time-lagged data were used for x(t)/x(t+7) **(A)** and x(t)/x(t21) **(B)**. White contour lines depict joint significance of periodicity. Black arrows reflect the phase difference in the areas with significantly joint periods. The solid red dots on the average power plot (the third from the left) indicate significantly joint periods at a probability of error 0.1. The color bars reveal the cross-wavelet power levels.

**Figure S3: Return Plots in 3 Dimensions for Poland.** New infections per day. **Top) Entire Observation Period.** 10^th^ March 2020 through 7^th^ November 2020. **Middle) Contained Phase.** Partial time frame through 18^th^ September 2020. **Bottom) Exacerbating Phase.** Partial time frame from 1^st^ September 2020.

**Figure S4: Bivariate wavelet analysis by US state.** The graphs display cross-wavelet power plot, wavelet coherence plot, average power plot and phase difference image (from left to right on each row) Time-lagged data were used for x(t)/x(t+7) **(A)** and x(t)/x(t21) **(B)**. White contour lines indicate significance of joint periodicity. Black arrows indicate the phase difference in the areas with significantly joint periods. The solid red dots on the average power plot (the third from the left) indicate significance at a level of 0.1.

**Figure S5: Evolution of Lyapunov exponents over time.** For a discrete mapping x(t+1) = F(x(t)) we calculate the local expansion of the flow by considering the difference of 2 trajectories. The Lyapunov characteristic exponent can be approximated as

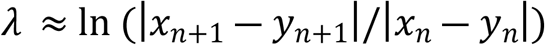

for 2 points *x_n_*,*y_n_* close to each other on the trajectory [https://www.math.tamu.edu/~mpilant/math614/Matlab/Lyapunov/LorenzSpectrum.pdf]. The changes of Lyapunov exponents are presented for the return plots of lags x(t+6) versus x(t), x(t+14) versus x(t), x(t+21) versus x(t), and x(t+35) versus x(t). **A) Countries.** Shown are ranges over 250 days. **B) US States.** Shown are ranges over 200 days. Mass. = Massachusetts.

